# Reconstitution of human fetal ovaries reveals niche requirements for primordial germ cell-like cell progression

**DOI:** 10.1101/2025.03.21.644608

**Authors:** Yolanda W. Chang, Marjolein Trimp, Talia Van Der Helm, Albert Blanch-Asensio, Arend W. Overeem, Susana M. Chuva De Sousa Lopes

## Abstract

Human primordial germ cell-like cells (hPGCLCs) can be specified from human induced pluripotent stem cells (hiPSCs), offering a valuable model for human germ cell development. However, further maturation steps of hPGCLCs rely on mouse feeders, or co-culture with mouse gonadal somatic cells. Exposure of hPGCLCs to human embryonic niche has not been attempted. Here, we co-cultured female hPGCLCs in two distinct somatic compartments. In reconstituted ovary (rOv) culture, human fetal germ cells proliferate and initiate meiosis, while hPGCLCs upregulate gonadal germ cell markers such as DDX4. Additionally, hPGCLCs can be supported in 3D culture by the amnion-like cells (AMLC) generated during PGCLC differentiation. Compared to rOV, hPGCLCs are less prone to dedifferentiation in PGCLC/AMLC aggregates. Finally, we demonstrated that SCF is crucial for the survival of hPGCLCs but not second trimester fetal germ cells. Together, this work highlights a shift in niche is required in human germ cell development.

**In Brief:** Chang and colleagues utilized in vitro reconstituted human fetal ovary (rOv) as somatic niche to mature human primordial germ cell-like cells (PGCLCs). hPGCLCs in rOv upregulate gonadal germ cell markers but are prone to dedifferentiation. In contrast, hPGCLCs cultured with amnion-like cells can be maintained without dedifferentiation. In both culture systems, SCF is crucial for the survival of hPGCLCs.

**Highlights:**

- Reconstituted human fetal ovaries (rOvs) support meiosis entry of fetal germ cells
- The rOVs support hPGCLCs to upregulate gonadal germ cell markers
- hPGCLCs show less dedifferentiation in amnion-like cell aggregates compare to rOv
- SCF is not required for survival of fetal germ cells, but crucial for hPGCLCs

## INTRODUCTION

Female fertility can be determined by many factors, but the most important is the size and quality of the oocyte reserve in the ovary, which is established during embryonic development. At around week 2 (W2) of development, a founder population known as primordial germ cells (PGCs) emerges in either the posterior epiblast or the amnion of the peri-implantation blastocyst (Hertig et al., 1958, Kobayashi and Surani, 2018). These PGCs, which give rise to the gametes, migrate through the hindgut endoderm epithelium and mesentery, reaching the genital ridge by W5 (Gomes Fernandes et al., 2018). Upon colonizing the developing gonad, PGCs upregulate gonadal germ cell markers, such as Deleted in Azoospermia Like (DAZL) and DEAD box protein 4 (DDX4) progressing to (pre-meiotic) oogonia (Nicholls et al., 2019, Guo et al., 2015). In females, from W11 onwards, some oogonia initiate meiosis in response to retinoic acid (RA) (Kurilo, 1981, Childs et al., 2011, Le Bouffant et al., 2010). These meiotic oogonia progress to bona fide oocytes by entering dormancy at the diplotene stage (dictyate arrest) and forming primordial follicles, structures that contain one oocyte associated with one layer of flat granulosa cells inside a surrounding basal membrane (Czukiewska and Chuva de Sousa Lopes, 2022). In vitro, human primordial germ cell-like cells (hPGCLCs) can be derived from human induced pluripotent stem cells (hiPSCs) (Sasaki et al., 2015, Irie et al., 2015). Recently, we also reported that hPGCLCs can be generated from hiPSCs alongside amnion-like cells using a basal membrane extract (BMEx) overlay culture (Overeem et al., 2023). Transcriptomics analysis revealed that hPGCLCs derived using these methods resemble pre-migratory PGCs, suggesting that differentiating hPGCLCs further to oocytes is still a significant challenge (Overeem et al., 2023, Irie et al., 2023). Additionally, isolated hPGCLCs are prone to de-differentiation during culture (Murase et al., 2020, Kobayashi et al., 2022). To propagate hPGCLCs while maintaining their germ cell identity, current approaches require culturing on feeder layers, coupled with the elimination of de-differentiated cells via FACS sorting, or the use of feeder-conditioned media (Murase et al., 2020, Kobayashi et al., 2022).

Yamashiro and colleagues have demonstrated that female hPGCLCs can differentiate into oogonia, able to upregulate DDX4 when aggregated with isolated somatic cells from embryonic day (E)12.5 mouse ovaries, forming reconstituted exogeneic rOvaries (xrOvaries), after co-culture for over 70 days (Yamashiro et al., 2018). This indicates the importance of the gonadal somatic environment for the maturation of hPGCLCs. However, it remains unknown whether hPGCLCs would undergo more efficient or further maturation when exposed to the human gonadal somatic niche, instead of the mouse gonadal somatic niche.

In this study, we developed a culture system using 3D reconstitution of dissociated second-trimester human fetal ovaries (rOv) to investigate the interaction between the female germ line and the female gonadal somatic niche. We demonstrated that the reconstituted gonadal somatic niche in rOv supports robust proliferation and meiosis initiation in fetal germ cells in culture using basal media alone. Furthermore, when hPGCLCs were introduced in the rOv, we observed upregulation of DDX4 and DAZL after 30 days of culture. hPGCLCs are known to de-differentiation after specification (Kobayashi et al., 2022, Murase et al., 2020), a phenomenon that we also observed when they are cultured within the fetal ovarian somatic niche. We found that co-culturing PGCLCs with amnion-like cells (AMLC) in 3D format allows PGCLCs to be maintained in culture for at least 2 weeks without showing de-differentiation. In conclusion, we demonstrated that the PGCLC/rOv and PGCLC/AMLC co-culture models are useful culture platforms to study human germ cell development.

## RESULTS

### Human fetal germ cells proliferate and enter meiosis in reconstituted ovary aggregates

To recreate the fetal ovary somatic niche in vitro, human fetal gonads from second trimester (W16, W17, W18, W19) were digested into single cells and allowed to reaggregate in either U or V-bottom low-attachment 96-well plates for 2 days to generate reconstituted ovary (rOv). The rOvs were then embedded in 1.5% agarose droplet and cultured further for more than 10 days (Figure 1A, 1B). Immunofluorescence staining of rOv sections revealed an increased number of POU5F1+ PGCs, POU5F1-DDX4+ premeiotic germ cells and SYCP3+ meiotic germ cells over time. The total germ cells increased from 1.9-7.9% on day 2 (D2), to 4.6-20% on day 12 (D12) (Figure 1C, D). Comparing the proliferating germ cells in W17 samples to d12 rOvs, we found that D12 rOvs have higher percentages of proliferating PGCs (Ki67+ and POU5F1+) and pre-meiotic germ cells (Ki67+ and DDX4+POU5F1-) (Figure S1A, B). To characterize the somatic cell types in the rOvs, we assessed the expression of FOXL2 and N2RF2, which mark (pre)granulosa and stromal cells respectively (Figure 1E). We observed that NR2F2+ stromal cells form the core of rOvs, whereas FOXL2+ granulosa cells and germ cells mostly localize in a peripheral ring. In resemblance of in vivo fetal ovary organization, (pre)granulosa cells (FOXL2+) tend to surround germ cells, while stromal cells (NR2F2+) cluster together and deposit collagen fibers (COL4) around them (Figure 1E). We also investigated whether agarose embedding is critical by culturing the rOvs in low attachment 96-well plate, and comparable results were observed in the rOvs cultured in suspension (Figure S1C, D). In summary, the rOv culture system recreates the germ cell niche in vitro which allows healthy proliferation of both germ cells and somatic cells.

**Figure 1.**
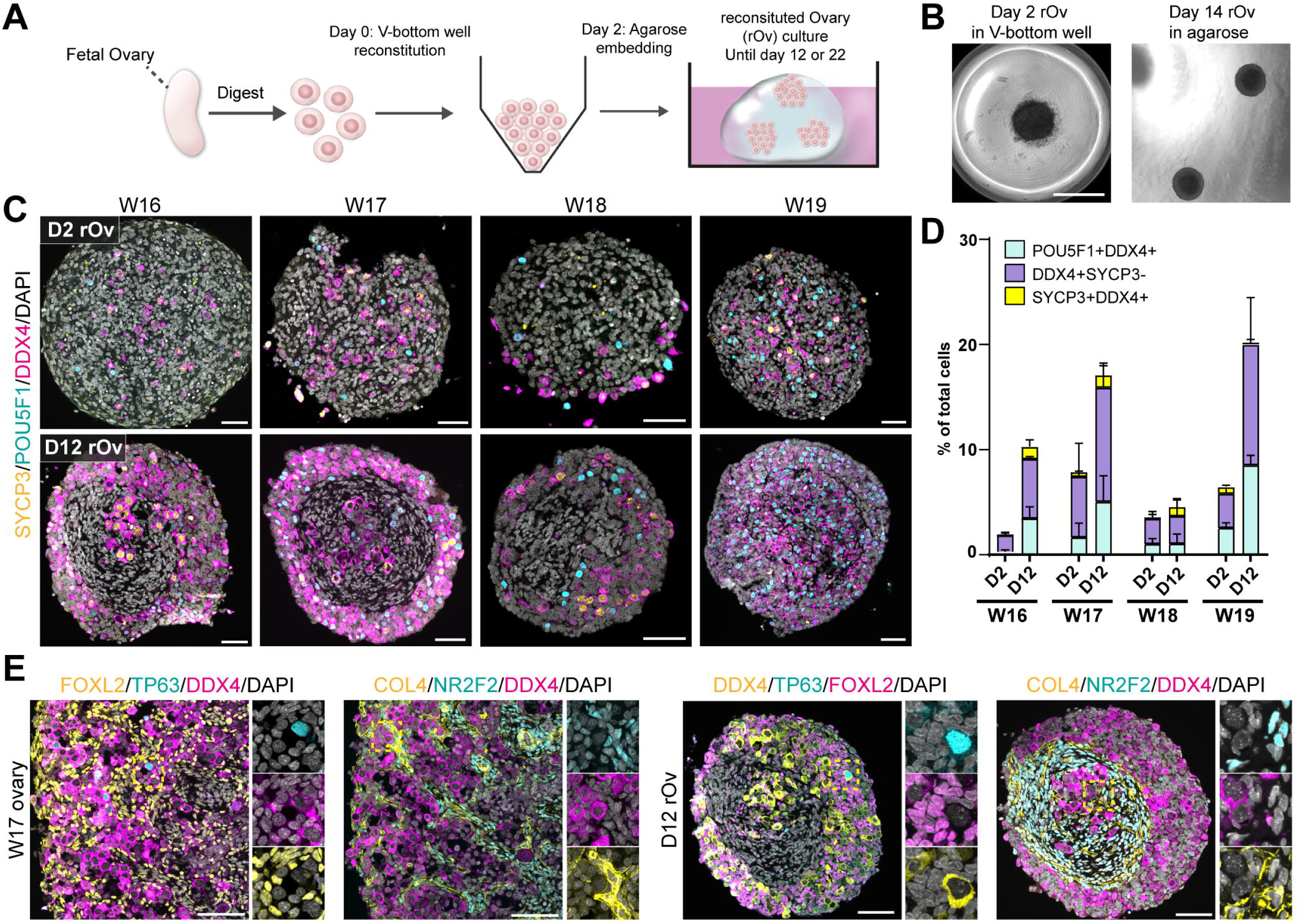
Maturation of fetal germ cells in the reconstituted fetal ovaries (rOv) **(A)** Schematics representing the aggregation of fetal ovary cells to form reconstituted ovaries (rOvs). **(B)** Bright field images of the rOvs at day 2(D2) and day 14 (D14) of culture. Scale bar: 750 μm. **(C)** Immunofluorescence (IF) images of the rOvs on D2 and D12, showing expression of POU5F1, DDX4 and SYCP3. Scale bar: 50μm. **(D)** Comparing percentages of PGCs (POU5F1+DDX4+), pre-meiotic germ cells (DDX4+SYCP3-) and meiotic germ cells (SYCP3+DDX4+) on D2 and D12, out of total number of cells per rOvs. Average of 2 to 4 rOvs were shown by column, error bars represent standard deviations (SD). **(E)** Representative IF images show expression of DDX4, FOXL2, TP63, COL4 and NR2F2 in both W17 ovary and reconstituted fetal rOvs. Scale bar: 50μm See also Figure S1.

### In vitro meiosis is promoted by retinoic acid signaling and progresses up to pachytene stage in rOv culture

Next, we examined the effects of adding SCF, Forskolin (FK), ascorbic acid (AA), BMP2, and RA on the proliferation and maturation of fetal germ cells in rOvs. SCF is known to be involved in PGC migration (Orth et al., 1997) and the survival of germ cells in the gonad (Dolci et al., 1991). Combining SCF with Forskolin increases the proliferation of both human and mouse PGCLCs (Ohta et al., 2017, Murase et al., 2020). Ascorbic acid acts as an antioxidant involved in gonadal tissue remodeling and apoptosis (Thomas et al., 2001). Retinoic acid, produced by the mesonephros, is essential for the initiation of meiosis (Bowles et al., 2006, Le Bouffant et al., 2010). BMP2 and RA together have been shown to induce mPGCLCs to enter meiosis prophase I in vitro (Miyauchi et al., 2017). On D12, we obtained similar percentages of PGCs, pre-meiotic germ cells, and total germ cells across all conditions (Figure 2A, S2A). However, the combination of BMP2, RA, and AA significantly improved the percentage of SYCP3+ cells compared to basal media (p=0.0453) (Figure 2A, S2A). By normalizing against the total number of germ cells, we found that SYCP3+ cells accounted for 9.5% of total germ cells in aRB27 (basal media) rOvs, and 25.4% in BMP2/RA/AA rOvs (Figure 2B). Culturing the W17 sample to D22 showed that the percentage of germ cells continue to increase across all conditions as reflected in immunofluorescence images and quantification (Figure 2C, D). More meiotic germ cells were found in both RA and BMP2/RA/AA conditions (24% and 21%) compared to other conditions (5-15%) (Figure 2D, S2B). Given that B27 supplement contains retinol, which can be converted to RA by enzymes present in the fetal gonad (Le Bouffant et al., 2010), we investigated whether fetal germ cells would enter meiosis without the presence of either retinol or RA. We observed the presence of meiotic germ cells in aRB27 without retinol or B27, although overall germ cell numbers were lower without B27 (Figure S2C).

**Figure 2.**
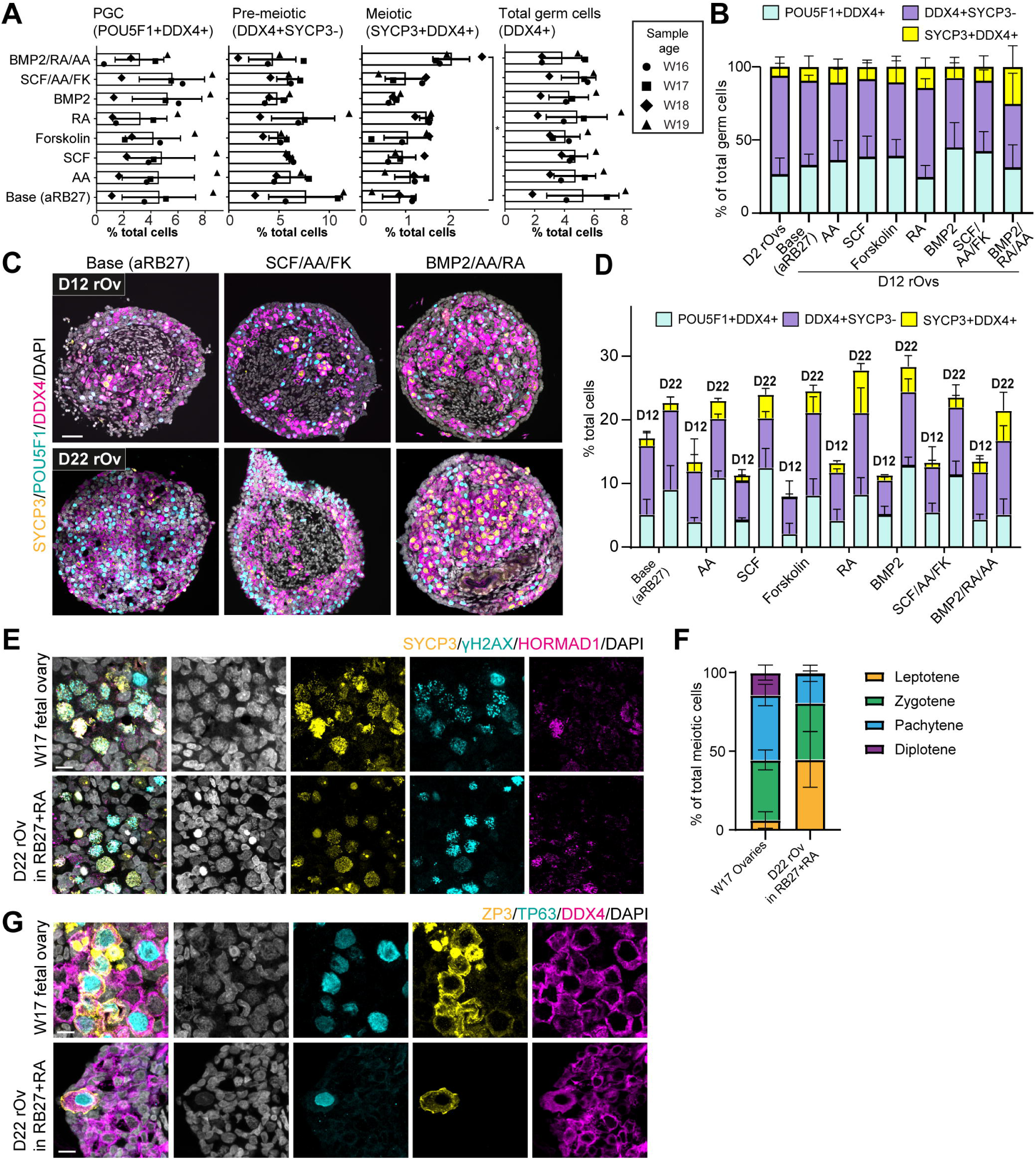
Effects of RA and SCF signaling on fetal germ cells in rOv (A) Average percentages of POU5F1+ cells, DDX4+SYCP3-cells and SYCP3+ cells normalized to total number of cells based on DAPI segmentation. Pooled from 4 different experiments using 4 different human fetal ovary samples (symbols indicate the week post fertilization ages). Individual points represent averaged values from each experiment. Error bars represent standard deviation, paired student t-test was performed between aRB27 and RA/BMP2/AA condition for % SYCP3+ Cells, *, p<0.05. (B) Normalizing the number of POU5F1+ cells, DDX4+SYCP3-cells and SYCP3+ cells from against total number of germ cells. Pooled from 4 samples, error bars represent SD. Representative IF images comparing the number of germ cells in D12 culture versus D22 culture. Scale bar: 50μm. (C) Comparing percentages of POU5F1+ cells DDX4+SYCP3- and SYCP3+DDX4+cells on D12 and D22 in the W17 sample (n=2-4) (D) IF images showing expression of SYCP3, γH2A.X and HORMAD1 in W17 fetal ovary and D22 reconstituted rOvs (cultured in aRB27+RA). Scale bar: 10μm (E) Percentage of cells in each meiotic prophase I stages based on expression of SYCP3, γH2A.X, HORMAD1 and DAPI staining. 4 fetal rOvs and W17 ovaries from 2 individuals were counted. (F) Representative images of diplotene cells expressing TP63, ZP3 and DDX4 in W17 fetal ovary and D22 reconstituted rOvs. Scale bar: 10μm See also Figure S2.

Meiosis prophase I can be divided into 4 stages based on the progression of homologous pairing and recombination: leptotene, zygotene, pachytene and diplotene. We compared the 4 stages of meiotic germ cells in rOvs to W17 fetal ovary based on the expression of meiotic markers and DAPI. Leptotene displays signs of chromosome condensation and expression of SYCP3, γH2AX and HORMAD1. Chromosome pairing begins in zygotene, as a results individual chromosome threads become more visible in DAPI, SYCP3 and γH2AX staining. Due to the synapsis between sister chromatids, pachytene is distinguished by concentrated focal spots of γH2AX and downregulation of HORMAD1. Diplotene oocyte does not express SYCP3 and are usually surrounded by a layer of granulosa cells (Zickler and Kleckner, 2023, Wang and Pepling, 2021). We found that majority of the meiotic germ cells in D22 rOvs from W17 sample are in leptotene (45%) and zygotene (36%) stages, only 19% are in pachytene. In W17 ovary, most are in zygotene (38%) and pachytene (41%) stages (Figure 2E, 2F). In W17 ovary, we observed around 15% of the meiotic germ cells to be in diplotene, but in cultured rOvs they are observed at very low frequency, marked by TP63 and ZP3 expression (Figure 2G, 2F).

SCF-KIT interaction plays an important role in anti-apoptosis and primordial follicle formation (Hoyer et al., 2005, Overeem et al., 2021), however we found adding recombinant SCF did not influence the percentages of germ cells in reconstituted fetal ovary (Figure 2A,B). Interestingly we observed a decrease of KIT, the SCF receptor, in rOv. In W17 fetal ovary, KIT is expressed on POU5F1+ PGCs, as well as POU5F1-DDX4+ germ cells but with lower intensity (Figure S2D). However, the expression of KIT is barely detectable in the rOv, in both aRB27 and SCF conditions (Figure S2D). Quantification of grey value per germ cells also showed drastic decrease of KIT (Figure S2E), suggesting the SCF-KIT interaction between germ cell and somatic cells might have been irreversibly disrupted, resulting in loss of KIT. To conclude, the reconstituted fetal ovary mimicked in vivo conditions, with fetal germ cells responding to RA and progressing into meiosis. However, meiotic progression might be slower in vitro as more leptotene cells and fewer diplotene oocytes were observed in rOvs compared to W17 ovary.

### The survival of hPGCLCs in reconstituted human fetal ovary is dependent on SCF

Next, we investigated whether the fetal germ cell somatic niche (rOv) can support the maturation of hPGCLCs. To do this we generated a hiPSC reporter line harboring POU5F1::GFP and DDX4::tdTomato (PG/DT) endogenous tags, allowing the tracing of iPSC derived germ cells from human germ cells (Figure 3SA). To reconstitute fetal ovary cells with hPGCLCs, PG/DT iPSCs were differentiated into PGCLCs using the 2D BMEx overlay method (Overeem et al., 2023). On Day 5 of differentiation, GFP+ITGA6+ PGCLCs were isolated by FACS sorting. For the hPGCLC reconstitution rOv, 5000 hPGCLCs were combined with 30,000 fetal ovary cells and allowed to aggregate in ultra-low attachment 96-well V-bottom well plate. After 2 days of aggregations (D2), hPGCLC containing rOvs were embedded in agarose droplet and cultured for up to 28 days (Figure 3A). On D2, GFP+ PGCLCs were detected in the rOvs by live cell imaging (Figure 3B). Flow cytometry analysis revealed that on D14, only 1.2% of the rOvs remained GFP+ under aRB27 conditions, while the addition of SCF increased the percentage of GFP+ cells to 2.3%. The percentage of GFP+ PGCLCs increased slightly on D21 and D30 in the SCF and SCF/AA/FK conditions but dropped below 1% in aRB27 (Figure 3C). Similar results were observed across different experiments (Figure 3D), demonstrating that SCF supplementation promotes the survival of hPGCLCs in the fetal ovary somatic niche. Live imaging of the rOvs on D29 confirmed higher percentages of GFP+ PGCLCs in the SCF and SCF/AA/FK conditions (Figure 3E). In the SCF/AA/FK condition, an increased amount of DDX4::tdTomato+ cells, also positive for germ cell markers TNAP and ITGA6, were observed (Figure 3F). This was further confirmed by colocalization of GFP (nuclei) and tdTomato (both nuclei and cytoplasm) signals in live-imaged rOvs (Figure 3G).

**Figure 3.**
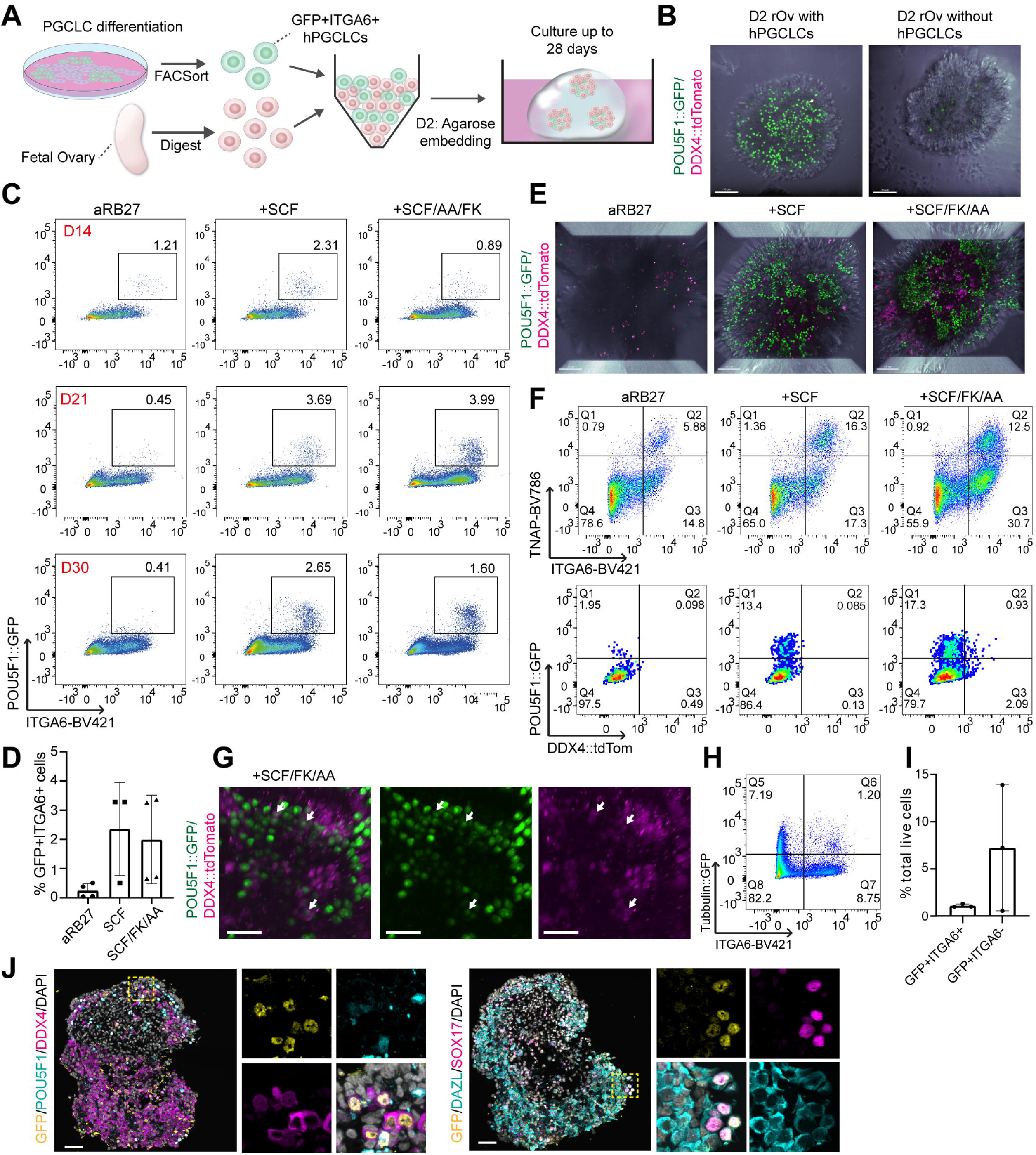
Maturation of hPGCLCs in the human fetal ovary niche (rOv) (A) Schematic representation of the reconstitution of PGCLCs with fetal ovary cells. (B) Live cell imaging showing POU5F1::GFP positive PGCLCs in D2 rOvs, in comparison to rOvs without PGCLCs as negative control. Scale bar: 100μm (C) FACS plot showing the percentage of POU5F1::GFP+ ITGA6+ hPGCLCs in reconstituted rOvs. (D) Bar graph depicting the average percentage of POU5F1::GFP+ ITGA6+ PGCLCs across four independent experiments (n=4). The rOvs were cultured for 21 to 25 days. Error bars represent SD. (E) Live cell imaging showing POU5F1::GFP and DDX4::tdTomato signals in D29 rOvs. Scale bar: 100μm (F) Top: FACS plots showing the all the germ cells positive for TNAP and ITGA6 in PGCLC/rOvs. Bottom: FACS plots showing cells positive for POU5F1::GFP and DDX4::tdTomato within the ITGA6+ TNAP+ population (Q2) (G) Zoomed in 3D live cell imaging showing the localization of POU5F1::GFP and DDX4::tdTomato signals, white arrows pointing at cells co-expressing GFP and tdTomato. Scale bar: 50μm (H, I) Percentages of TUBA1B::GFP+ PGCLCs (ITGA6+) and TUBA1B::GFP+ differentiated cells (ITGA6-) measured by FACS, n=3, error bars represent SD. (J) Representative IF images showing TUBA1B::GFP+ cells expressing POU5F1, SOX17, DDX4 and DAZL. Scale bar: 50μm. See also Figure S3.

When 5,000 hPGCLCs were added to 30,000 fetal ovary cells to rOvs, hPGCLCs initially comprised 14% of the rOvs at plating. However, by Day 30, less than 3% of the cells remained as hPGCLCs. This prompted us to investigate whether the hPGCLCs were outcompeted by fetal ovary cells or had differentiated into other cell types. To trace all hiPSC-derived cells during co-culture, we generated TUBA1B –tagged hiPSC lines (TUBA1b::GFP and TUBA1B::Scarlet) (Figure S3A). Once again, 5000 Tubulin-GFP+ ITGA6+ hPGCLCs were isolated and reconstituted with 30,000 fetal ovary cells and cultured using the same method. We observed that only 1.2% of total cells are hiPSC-derived germ cells positive for both GFP and ITGA6. However, the majority of hiPSC-derived cells (around 7% of cells in the rOvs), were TUBA1B::GFP+ but negative for ITGA6, indicating the differentiation of TUBA1B::GFP+ cells into non-PGCLCs (Figure 3H, 3I). This effect was greatly exacerbated by the addition of RA, leading to 79% Scarlet+ ITGA6-cells without SCF/AA/FK and 55% with, indicating a loss of germ cell fate and the proliferation of unknown somatic cell types (Figure S3B). Lastly, immunofluorescence confirmed that TUBA1B-GFP+ cells that are still PGCLCs, indeed express PGC markers such as POU5F1 and SOX17, some of them also upregulated DAZL and DDX4 (Figure 3J). To summarize, we demonstrated that the rOvs can support the maturation of hPGCLCs to upregulate DDX4 and DAZL. However, majority of hPGCLCs in the rOv struggle to maintain germ cell fate, which may contribute to their limited upregulation of DDX4. Additionally, unlike fetal germ cells, PGCLCs rely on SCF supplementation for survival in the rOvs.

### PGCLC/AMLC 3D aggregate culture allows long-term maintenance of hPGCLCs

hPGCLCs were shown to resemble pre-migratory PGCs based on transcriptomic studies (Overeem et al., 2023), it is possible that the second trimester somatic niche are “too mature” for the PGCLCs. When PGCLCs are induced from iPSCs, they are specified alongside AMLC, mimicking the in vivo environment of nascent PGCs. We hypothesized that hPGCLCs would have a lower tendency to differentiate into other cell types when co-cultured with AMLCs. To test this, hiPSCs were differentiated into hPGCLCs, on day 5, the culture was dissociated into single cell suspension. Aggregates of 3000 cells were formed using Aggrewell microwell plates and transferred into Vitrogel for further culturing (Figure 4A, 4B). We also tested whether adding SCF, AA and FK could affect the survival of PGCLCs.

**Figure 4.**
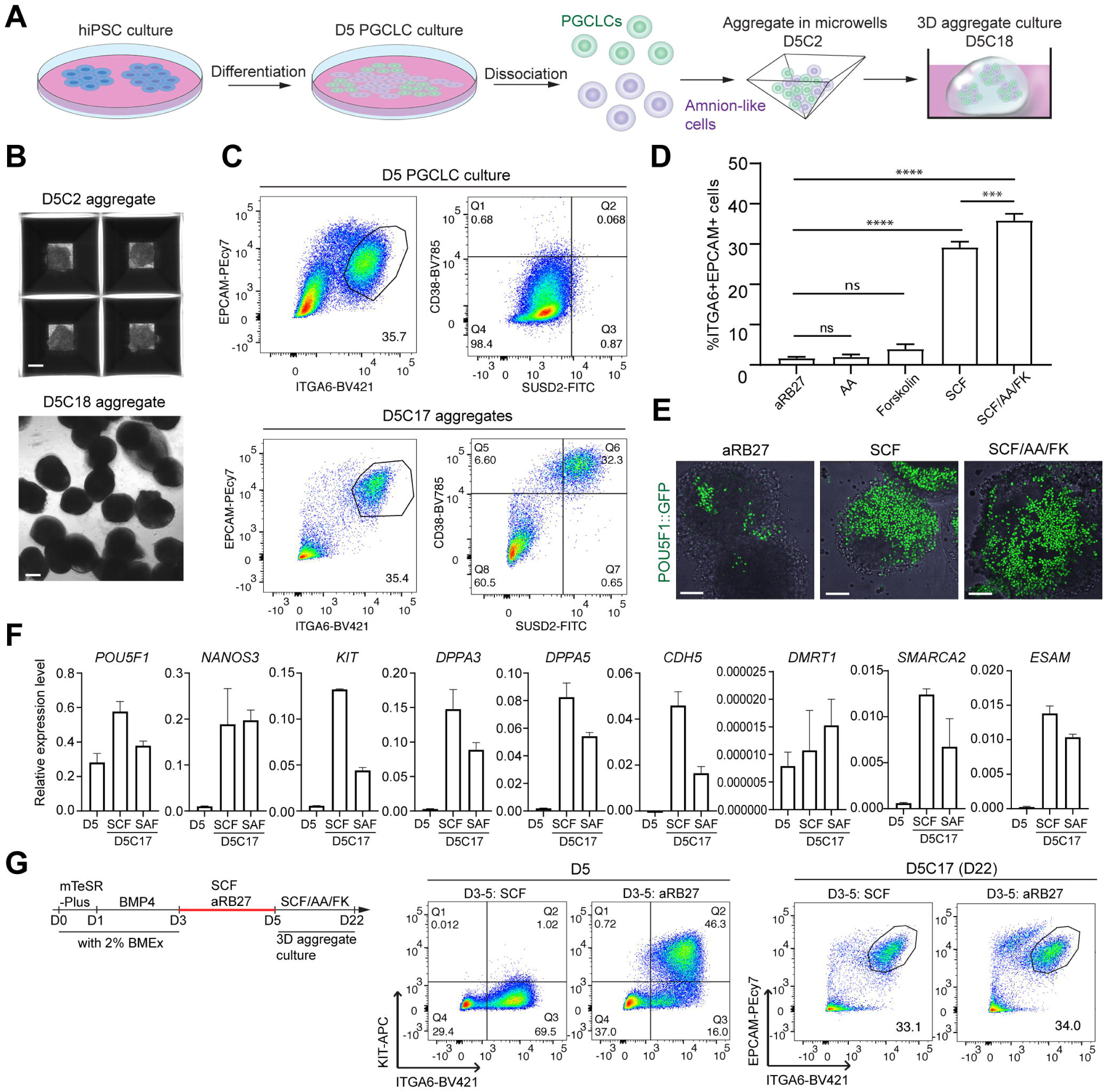
Establishment of PGCLC/AMLC aggregate culture for the maintenance of PGCLCs (A) Schematic representation of the aggregation and culture of PGCLC/amnion like-cell (AMLC) aggregates. (B) Brightfield images showing the aggregates on day 2 (D5C2) in Aggrewell, and on day 18 (D5C18) in VItrogel. Scale bar: 150μm (C) FACS plots showing expression of EPCAM, ITGA6, SUSD2 and CD38 in D5 PGCLC differentiation culture and D5C17 PGCLC aggregates. Aggregates were cultured in SCF/AA/FK condition. (D) Bar graph showing the percentage of ITGA6+EPCAM+ cells in PGCLC/amnion-like cells aggregates. n=3, error bar depicts SD. One-way ANOVA analysis: p<0.0001. * represents level of significance for individual comparisons done based on Tukey multiple comparison test. ****: p<0.0001, ***: p<0.001 (E) Live cell 3D imaging showing POU5F1::GFP+ PGCLCs in D5C17 aggregate. Scale bar: 100μm (F) Relative gene expression level of indicated genes normalized to *GAPDH*, assessed by real-time qPCR. n=3 technical replicates, error bars represent SD. (G) Testing the effect of SCF treatment during PGCLC differentiation. Left: schematic of experimental design, SCF is either added or not added during D3-5 of differentiation. Center: FACS plots showing the upregulation of KIT at D5 without SCF. Right: FACS plot showing the percentage of EPCAM+ITGA6+PGCLCs in D5C17 aggregates. See also Figure S4.

After culturing for 17 days (D5C17), PGCLC/amnion-like cell aggregates were able to maintain around 35% PGCLCs in the SCF/AA/FK condition, consistent with the initial 35% at the start of the aggregate culture (Figure 4C). Comparing to without SCF, barely any PGCLCs were left in the aggregates, demonstrating that PGCLC survival depends on SCF. Although adding Forskolin slightly increased the survival of hPGCLCs slightly to 3.9% (ns), combining SCF and Forskolin further improved the percentage of hPGCLCs compared to SCF (Figure 4D).

Using the PG/DT hiPSC lines, we were able to visualize the POU5F1::GFP+ hPGCLCs in the aggregates using 3D live cell imaging (Figure 4E). Although we could not detect expression of DDX4 (Figure S4A), we observed upregulation of PGC markers CD38 and SUSD2, which are not expressed on D5 PGCLCs (Figure 4C). Furthermore, PGCLCs in the aggregate culture also upregulated expression of PGC markers such as KIT, DPPA3, DPPA5, SMARCA2 and ESAM. DMRT1 and CDH5 were previously described to be upregulated in migratory PGCs (Irie et al., 2023), we found that CDH5 was upregulated in D5C17 PGCLC aggregates, but not DMRT1 (Figure 4F). Interestingly we could not detect the expression of KIT on PGCLCs by flow cytometry (Figure S4B), even though hPGCLCs depends on its ligand SCF strongly for its survival. Others have reported that using soluble SCF as KIT ligand can lead to the internalization of KIT (Chen et al., 2017a, Miyazawa et al., 1995), we tested if we can promote the expression of KIT by omitting SCF during the differentiation (Figure S4C). We observed that removing BMP4, LIF and EGF during D3-5 do not affect the efficiency of the differentiation, while omitting SCF leads to 10% decrease in hPGCLC differentiation efficiency (Figure S4C). However, not adding SCF results in drastic upregulation of KIT detected by flow cytometry (Figure 4G) and immunostaining (Figure S4D). Furthermore, we investigated whether the transcript expression of KIT is affected by adding SCF, we saw that in the aRB27 condition, KIT is expressed 3 folds higher than in the SCF condition (Figure S4E). Nevertheless, this difference in KIT expression does not affect the survival of hPGCLCs in 3D PGCLC aggregates. After 17 days of culture in SCF/AA/FK media (D5C17), both aRB27 and SCF (D3-5) conditions yield similar percentages of PGCLCs (Figure 4G). In conclusion, we have developed a 3D model that maintains the germ cell fate of hPGCLCS for over two weeks, without the need of isolating hPGCLCs from its differentiation culture. Although co-culturing hPGCLCs with AMLCs did not lead to DDX4 upregulation, it better supports the maintenance of hPGCLCs compared to PGCLC-rOv with less dedifferentiated cells. Our two in vitro models highlighted the importance of the somatic niche in maintaining the cell fate of hPGCLCs.

## DISCUSSION

In this study, we successfully reconstructed the fetal ovary somatic niche in vitro using human fetal ovary cells from the second trimester. Both fetal germ cells and (pre)granulosa cells proliferated in the base aRB27 medium without the need for additional growth factors, demonstrating that the presence of a somatic cell niche alone is sufficient for germ cell growth. The rOvs closely mimicked in vivo conditions, with fetal germ cells responding to RA and progressing into meiosis. However, compared to the W17 ovary, rOVs exhibited significantly more leptotene cells and fewer diplotene oocytes, suggesting that meiotic progression may be slower in rOVs. We also found that while exogenous RA increases overall meiosis entry, it is not essential, confirming previous findings that RA is synthesized by the gonad in humans (Le Bouffant et al., 2010). Remarkably, meiosis initiation still occurred in rOVs even without retinol, indicating that constant endogenous RA production might not be necessary. Together, our results demonstrated that human meiosis initiation might be intrinsic to the gonads, does not dependent on RA signaling from the mesonephros as in mice (Bowles et al., 2006). Unlike fetal germ cells, addition of RA induced PGCLC to dedifferentiate into other cell type.

We also demonstrated that the somatic cells from the D5 hPGCLC differentiation culture can be utilized as a niche to support the in vitro maintenance of hPGCLCs. This approach eliminates the need for laborious isolation of hPGCLCs via FACS sorting, making the AMLC/PGCLC 3D aggregate culture a valuable model for studying long-term cultured PGCLCs. By comparing the co-culture of hPGCLCs with two different somatic cell types, we highlighted how distinct microenvironments influence hPGCLCs’ fate decisions, whether to maintain their germ cell identity, undergo maturation, or differentiate into other cell types. Although PGCLCs can survive and maintain their PGC identity better when co-cultured with AMLCs, we did not observe upregulation of DAZL. This finding aligns with previous research suggesting that signaling from the gonad or genital ridge is necessary for germ cells to upregulate DAZL (Hu et al., 2015). Interestingly, co-culturing with amnion-like cells led to the upregulation of SUSD2 and CDH5 which are markers of migratory PGCs (Irie et al., 2023, Hwang et al., 2020).

Using our two in vitro culture models, we have gained more insights into the role of SCF in germ cell development. SCF is crucial for the survival of mouse PGCs (mPGCs) during migration in vivo (McCoshen and McCallion, 1975), as well as in culture (Dolci et al., 1991). In human second trimester fetal ovary, SCF is expressed by the (pre)granulosa cells and KIT is expressed by fetal PGCs and mitotic germ cells (Overeem et al., 2021). The SCF/KIT interaction is crucial for primordial follicle formation in the fetal ovary (Jin et al., 2005, Jones and Pepling, 2013). To our surprise, SCF did not improve the proliferation of human fetal germ cells. Additionally, KIT is significantly downregulated, indicating a loss of SCF/KIT interaction in rOVs. The low number of diplotene oocyte in rOvs may be related to this loss of interaction, although supplementing SCF did not appear to affect the survival of PGCs, pre-meiotic, or meiotic germ cells. Since KIT has been observed to downregulate in meiotic germ cells (Overeem et al., 2021, Hoyer et al., 2005), this suggests SCF might not be necessary for the survival of fetal PGCs and pre-meiotic germ cells, but crucial for the survival of primordial follicles.

Despite the lack of effect on fetal germ cell proliferation, SCF significantly improved the survival of PGCLCs in the rOvs and PGCLC/AMLC aggregates. Since PGCLCs resemble week 3-4 PGCs, this indicates that soluble SCF supplementation is more crucial for the survival of early PGCs than late. The receptor of SCF, KIT, is a marker for both migratory and gonadal germ cells. We found KIT to be absent on hPGCLCs but can be induced by removing SCF from the differentiation protocol. Omitting SCF for just two days in the differentiation resulted in approximately 10% decrease in differentiation efficiency, highlighting once more the importance of SCF in hPGCLC survival. Interestingly omitting LIF or EGF did not influence the differentiation efficiency. The loss of KIT in culture could be a results of receptor internalization in response to soluble SCF (Miyazawa et al., 1995, Chen et al., 2017a). Remarkably, the lack of KIT expression in newly specified D5 hPGCLCs did not impair their ability to respond to SCF and maintain its germ cell fate in long-term culture. In summary, we have established two in vitro models for the maturation and maintenance of hPGCLCs in human somatic niche. Using these models, we elucidated the role of SCF in female germ cell development in vitro.

### Limitation of the study

Migratory PGCs reach the developing gonad at W5-6, we speculated that second trimester fetal ovary might be too mature for hPGCLCs, it would be more appropriate to reconstitute hPGCLCs with W5-6 human fetal ovaries which we rarely have access to. Nevertheless, it has been reported that when PGCLCs were co-cultured with mouse E12.5 fetal ovary somatic cells (equivalent of human W7), dedifferentiation of hPGCLCs was also observed (Yamashiro et al., 2020).

## AUTHOR CONTRIBUTIONS

Conceptualization: Y.W.C., A.W.O., S.M.C.d.S.L.; Methodology: Y.W.C., M.T., T.V.D.H., A.B-A., A.W.O., S.M.C.d.S.L.; Investigation: Y.W.C., M.T., T.V.D.H., A.B-A., A.W.O., S.M.C.d.S.L.; Formal Analysis: Y.W.C., A.W.O., S.M.C.d.S.L.; Writing: Y.W.C., M.T., T.V.D.H., A.B-A., A.W.O., S.M.C.d.S.L.; Resources: M.T., T.V.D.H., S.M.C.d.S.L.; Funding Acquisition: S.M.C.d.S.L.; Supervision: S.M.C.d.S.L. All authors approved the final version of the manuscript.

## ACKNOWLEDGEMENTS

We would like to thank the patients who donated the human fetal material used in this study as well as the staff of the Gynaikon Clinic in Rotterdam and het Vrelingshuis in Utrecht. We thank Dr. K. Szuhai, Dr. C. Freund, Dr. R. Davis and members of the Chuva de Sousa Lopes group for fruitful discussions, and S.Hillenius for technical assistance. This work was supported by the Dutch Research Council (VICI-2018-91819642 to Y.W.C., A.W.O., and S.M.C.d.S.L.), the Nederlandse organisatie voor gezondheidsonderzoek en zorginnovatie (ZonMw) (PSIDER-10250022120001 to T.V.D.H. and S.M.C.D.E.S.L.), and the Novo Nordisk Foundation (reNEW NNF21CC0073729 to A.W.O., M.T., and S.M.C.d.S.L.).

## Data availability

This study includes no data deposited in external repositories.

## METHODS AND MATERIALS

### Human samples and ethics statement

All experiments conducted in this study followed the guidelines specified in the Declaration of Helsinki for Medical Research involving Human Subjects. For ethics approval, a letter of no objection was issued by the Medical Ethical Committee of Leiden University Medical Center (B21.054).

The human fetal ovary samples used in this study were collected from elective abortions without medical indication, after obtaining informed consent from the donors. Human fetal ovaries were dissected and rinsed in 0.9% NaCl solution (Fresenius Kabi). To cryopreserve the fetal ovary, the whole ovary was first incubated in 0.25% Trypsin-EDTA (Thermo Fisher Scientific) at 37 °C for 30 minutes, with radial rotation at 250rpm. The trypsin was then removed, and the fetal ovary was washed with 1:1 mix of low glucose DMEM (Thermo Fisher Scientific) and M199 media (Thermo Fisher Scientific). The ovary was then dissected into pieces of roughly 1mm and slow-frozen in Bambanker (Nippon Genetics).

### Fetal ovary reconstitution

To create the reconstituted fetal ovary aggregates (rOv), frozen fetal ovary samples described from the previous section are thawed at 37 °C and rinsed with DMEM:F12 media (Thermo Fisher Scientific). Fetal ovary pieces were digested as previously described (Overeem et al., 2023). Briefly, the ovary pieces were incubated with 1mg/ml Collagenase IV (Invitrogen), 0.5mg/ml Hyaluronidase (Sigma-Aldrich) and 20U/mL DNase I (Sigma-Aldrich) at 37 °C for 30-45 minutes, followed by vigorous pipetting and filtering through a 40 µm nylon mesh strainer (Corning). For the fetal ovary cells to aggregate, 30,000 fetal ovary cells, either with or without 5000 FACS-sorted PGCLCs were incubated in aRB27 basal medium [advanced RPMI1640 (Thermo Fisher Scientific) supplemented with B27 (1:100) (Thermo Fisher Scientific),Glutamax (Thermo Fisher Scientific), MEM Non Essential AminoAcids (Thermo Fisher Scientific) and Mycozap(Lonza)] plus RevitaCell supplement (Thermo Fisher Scientific), in 96-well ultra-low attachment V-bottom and U-bottom well plates (S-bio). For the floating culture condition, rOvs were left in the ultra-low attachment well for culturing, with media change every 3 days. For rOvs cultured in agarose, the rOvs were embedded in 1.5% low-melting point agarose (Promega) droplet in aRB27 media with the addition of ascorbic acid (Sigma-Aldrich), SCF (R&D Systems), forskolin (Biogems), BMP2 (R&D Systems), and retinoid acid (Sigma-Aldrich), as specified in the figures and results.

### Generation of hiPSC transgenic lines

LUMC0099iCTRL04 iPSC line (iPSC reg LUMCi004-A, characterizations can be found: https://hpscreg.eu/cell-line/LUMCi004-A) was cultured in mTeSR-Plus media (STEMCELL Technologies) supplemented with MycoZap (Lonza), on tissue culture plates coated with Geltrex (Thermo Fisher Scientific) diluted in DMEM/F12 (Thermo Fisher Scientific) at 1% (v/v) concentration. Cells were cultured at 37 °C in a humidified normoxic incubator with 5% CO2. Routine clump passaging was performed every 4-7 days using ReLeSR at 1:5-1:30 split ratio (Stem Cell Technologies). Routine mycoplasma testing was performed.

For generation of endogenous reporter lines, knock-in vectors were constructed by molecular cloning. All PCR-amplifications were performed using PrimeSTAR MAX (Takara) according to manufacturer’s instructions. For generation of a POU5F1-T2A-GFP-NLS knock-in vector, homology arms, EGFP and the plasmid backbone were PCR-amplified from the AX74_pDonorOCT4.TS knock-in vector (Chen et al., 2020). The T2A sequence was included as an overhang sequence during PCR. SV40 NLS sequence was generated by annealing of sense and antisense oligonucleotides. These sequences were then assembled as depicted in Figure S3A, using NEBuilder® HiFi DNA Assembly Master Mix (New England Biolabs) according to manufacturer’s instructions. For Scarlet-TUBA1B knock-in vector, homology arm sequences flanking the start codon at the N-terminus, as well as FLAG-Scarlet were ordered as synthesized products (Baseclear) and assembled using NEBuilder® HiFi DNA Assembly Master Mix. For generation of DDX4-tdTomato and TUBA1B-3xEGFP knock-in vectors, homology arm sequences at the C-terminus of these genes were PCR-amplified from genomic DNA, with overhangs for subsequent NEBuilder® HiFi DNA Assembly (See Table S2 for primer sequences). Genomic DNA was isolated from hiPSC line F99 using the DNeasy Blood & Tissue Kit (Qiagen). EGFP, T2A and SV40-NLS sequences were obtained as described for the POU5F1-T2A-GFP-NLS vector, tdTomato was PCR-amplified from Bxb1-tdTomato plasmid (Addgene, #198041) (Blanch-Asensio et al., 2024), and a loxp-flanked PGK-PuroR cassette was ordered as synthesized product (Baseclear). Various parts were then merged as depicted in Figure S3A, using NEBuilder® HiFi DNA Assembly, in a linearized pUC57-KAN (for DDX4) or pUC19 (for TUBA1B) backbone vector. Guide RNAs targeting TUBA1B N-terminus, TUBA1B C-terminus and DDX4 were inserted into the pAY56 gRNA gRNA spacer acceptor construct AY56_pU6.opt-sgRNA.BveI-stuffer (Maggio et al., 2020) by oligo annealing and ligation using T4 DNA Ligase (New England Biolabs). For POU5F1 targeting, in trans paired nicking (Chen et al., 2017b) was used by transfecting AX33_pgRNAOCT4.1 (Chen et al., 2020), together with AB65 pCAG.Cas9D10A.rBGpA and HDR donor plasmid POU5F1-T2A-GFP-NLS. This DSB-independent gene editing method allows for generating cell populations and organoids with precise gene knock-ins without concomitant target allele disruptions (Bollen et al., 2022, Wang et al., 2023).

To generate knock-in LUMC0099iCTRL04 lines, 100,000 cells were plated on a well of 12-well plate, 24 hours later, transfected with HDR donor plasmids, pAY56 based gRNA expression plasmids, and Cas9-D10A nickase expression plasmid AB65_pCAG.Cas9D10A.rBGpA (Chen et al., 2020) using Lipofectamine™ Stem Transfection Reagent (Thermo-Scientific) following manufacturer’s instructions. In case of TUBA1B and POU5F1, transfected cells were passaged two days after transfection, expanded for five days, and GFP-positive cells were then isolated by FACS. In case of DDX4, 72 hours after transfection, cells were treated with 0.5 ug/ml puromycin (Invitrogen) for 72 hours. Surviving clones were expanded, passaged, and treated with TAT-CRE Recombinase (Merck-Millipore) according to manufacturer’s instructions. After 48 hours, Cre-treated cells were passaged for clonal dilution, and resulting expanded clones were screened for successful tdTomato insertion, and PGK-PuroR cassette removal by PCR.

### 2D hPGCLC differentiation

HiPSCs were differentiated into PGCLCs as described previously (Overeem et al., 2023). Briefly, single cell suspension of hiPSCs were plated on Geltrex coated plates in mTeSR-Plus medium containing RevitaCell and 2% Geltrex. On day 1 and day 2 after plating, the media was changed to aRB27 containing BMP4 and 2% Geltrex. On day 3 and day 4, the media was switched to aRB27 containing 50ng/ml SCF or indicated otherwise. Day 5 PGCLC cultures were dissociated by incubating with Accutase at 37 °C for 15-20 minutes and pipetting up and down to break clumps into single cells for downstream use.

### Culturing PGCLCs in 3D aggregate with amnion-like cells

Day 5 PGCLC culture were dissociated into single cell suspension as described previously, 9 x 10^5^ cells were added to a well of Aggrewell 800 (24-well format) (Stemcell Technologies) in mTeSR-Plus media supplemented with RevitaCell. The plate was centrifuged at 100g for 3 minutes, then allowed for the aggregates to form for 2 days. Rest of the D5 PGCLCs were analyzed with flow cytometry for PGCLC percentage. On day 2, the aggregates were collected and washed with aRB27 media; then 1/10 of the total aggregates were embedded in 50µL droplets of 50% diluted 3D high concentration Vitrogel (diluted with Vitrogel dilution solution Type I)(The Well Bioscience). The gel droplets were allowed to solidify at 37 °C for 15 minutes. Then aRB27 media supplemented with ascorbic acid, SCF, or forskolin are added gently to the well. Media exchange was performed every 2-3 days for up to 2 weeks. On Day 17, the aggregates were recovered by rinsing with DPBS-/- (Thermo Fisher Scientific) and mechanically breaking the gel.

### Flow cytometry and fluorescence-activated cell sorting (FACS)

For fluorescence-activated cell sorting of PGCLCs, single cell suspension of Day 5 PGCLCs were filtered through a 40µM nylon mesh strainer twice (Corning), washed once in FACS buffer [DPBS with 0.5% BSA(Sigma-Aldrich)] and resuspended with conjugated-antibodies diluted in FACS buffer. The cells were incubated together with the antibodies for 30 minutes on ice and kept in the dark. Thereafter, the cells were washed once by centrifugation and resuspended in FACS buffer containing 7AAD(BioLegend,1:100). A CytoFLEX SRT benchtop cell sorter with 100µm nozzle (Beckman) was used to sort the cells.

For Flow cytometry analysis of rOvs and PGCLC/amnion-like cell aggregates, the tissues were removed from the gel droplets and digested with 10mg/ml collagenase IV (Invitrogen) for 30-60 minutes at 37 °C with pipetting in between. Once the tissues became single cells, the cells were filtered through a 35 µm nylon mesh strainer (Falcon), stained with conjugated antibodies and washed the same way as described above. The flow cytometry analysis was performed on an LSR Fortessa flow cytometer (BD biosciences) and FlowJo Software (BD Biosciences) were used for analysis.

### Immunofluorescence and Histology

Human fetal ovaries and fetal rOvs were fixed in 4% PFA at 4°C overnight, then rinsed 3 times with PBS. Due to the small size of the rOvs, they were then embedded in 4% Agarose droplet. For paraffin embedding, fetal ovary samples and fetal rOvs were embedded with a Shandon Excelsior tissue processor (Thermo Fisher Scientific). The samples were then sectioned at 5µm thickness using an RM2065 microtome (Leica Instruments) and mounted on Star Frost glass slides (Knittel). For immunofluorescence, sections were deparaffinized in Xylene and subsequently dehydrated with 100%, 90%, 80%, 70% ethanol ending in water. Antigen retrieval was performed by incubating the slides in TRIS-EDTA buffer (10 mM Tris, 1 mM EDTA, 0.05% Tween 20, pH 9) at 98 °C for 12 minutes in a TissueWave 2 microwave (Thermo Fisher Scientific). An extra antigen retrieval step was added forGFP stainings. In this case, the slides were incubated with 1%SDS for 5 minutes at room temperature(rt). The treated slides were then washed in PBS two times and blocked with blocking solution [2% BSA and 10% normal donkey serum in PBST (0.05% Tween 20 in PBS)]. When staining for SYCP3, 0.5% Triton was also added to the blocking solution. After blocking at rt for 1 h, primary antibody slides were then incubated with primary antibodies diluted in blocking solution overnight at 4°C in a humidified chamber. When staining with anti-GFP antibody, slides were incubated at 37°C overnight. The slides were then washed three times in PBST, incubated with secondary antibodies and DAPI (Sigma-Aldrich) diluted in PBST with 2% BSA at rt for 1h. The samples are then washed three times in PBS, mounted with coverslips in ProLong Gold (Thermo Fisher Scientific), and allowed to dry for 24 hours in the dark before imaging.

### Imaging and quantification

Both live cell imaging and paraffin sections were imaged on either 200 or 500 series Dragonfly spinning disk confocal microscope (Andor). Images were processed either in Fiji (ImageJ2) or Imaris. Quantification of the PGC, oogonia and meiotic germ cells were done by manual counting of the largest cross section of the rOv, then normalizing the germ cell number by total cell number in the same section. Total cell number of a section was quantified based on DAPI segmentation using Stardist (Caicedo et al., 2019). 1-4 rOvs were quantified per condition. For KIT expression, the tissue sections were stained with primary and secondary antibodies at the same time, imaged at the same time using same settings. KIT signals were quantified using “Plot Profile” of imageJ, background value was subtracted and area under the curve was calculated for the total grey value. Finally average grey value per germ cell was normalized by the number of germ cells.

### RNA isolation and quantitative RT-PCR

To isolate total RNA, D5 or D5C17 culture were first digested into single cell suspension as described in previous sections, followed by centrifugation to form a pellet. RNA was then extracted from the pellet using RNeasy Micro Kit (Qiagen). cDNA was synthesized using iScript-cDNA Synthesis kit (Bio-Rad). For the real-time PCR, 10ng cDNA product was added to a 10ul qPCR reaction, with iTaq Universal SYBR Green Supermixes (Bio-Rad) and 250nM forward and reverse primers (primers used can be found in Supplemental Table 2). The PCR reactions were run on a BioRad CFX384 Real Time PCR System. To calculate the relative gene expression level (2^(-delta Ct)), delta Ct was calculated by normalizing to housekeeping gene *GAPDH* or *HPRT1*.

### Statistical analysis

Column plots were generated using either GraphPad Prism 9 software or numpy and matplotlib packages. Statistical analyses were performed using GraphPad Prism 9. Specifically, for Figure 2A paired student t-test was used to compare % meiotic germ cells in aRB27 with BMP/RA/AA conditions. For Figure 4D, first the data was confirmed to have normal distribution with Shapiro-wilk test, then One-way ANOVA with Tukey’s multiple comparison test was used for multiple pair-wise comparisons.

