## Supplemental figures and tables for "Reconstitution of human fetal ovaries reveals niche requirements for primordial germ cell-like cell progression"

Supplementary

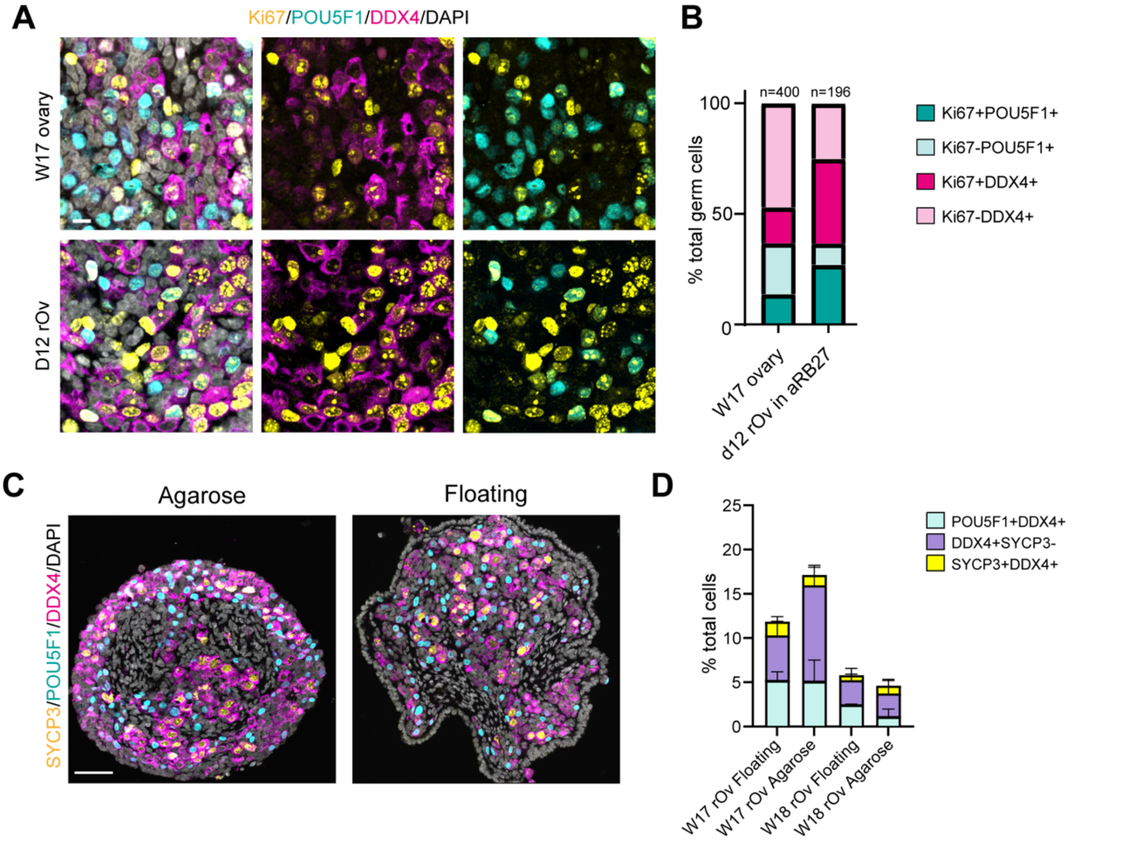


**Supplementary Figure S1. Human fetal ovary reconstitution**

(A) Representative IF images showing expression of Ki67, POU5F1 and DDX4 in both W17 fetal ovary and reconstituted fetal rOvs. Scale bar: 10μm.

(B) Quantification of proliferating primordial germ cells (POU5F1+) and pre-meiotic germ cells (POU5F1- DDX4+) in both W17 fetal ovary and rOvs. N represents number of germ cells counted.

(C) Representative IF images of D12 rOvs cultured in suspension (floating) or embedded in agarose. Scale bar: 50μm.

(D) Average percentages of POU5F1+ cells, DDX4+SYCP3- cells and SYCP3+ cells in floating and agarose condition, 2-3 rOvs per condition were used for quantification.

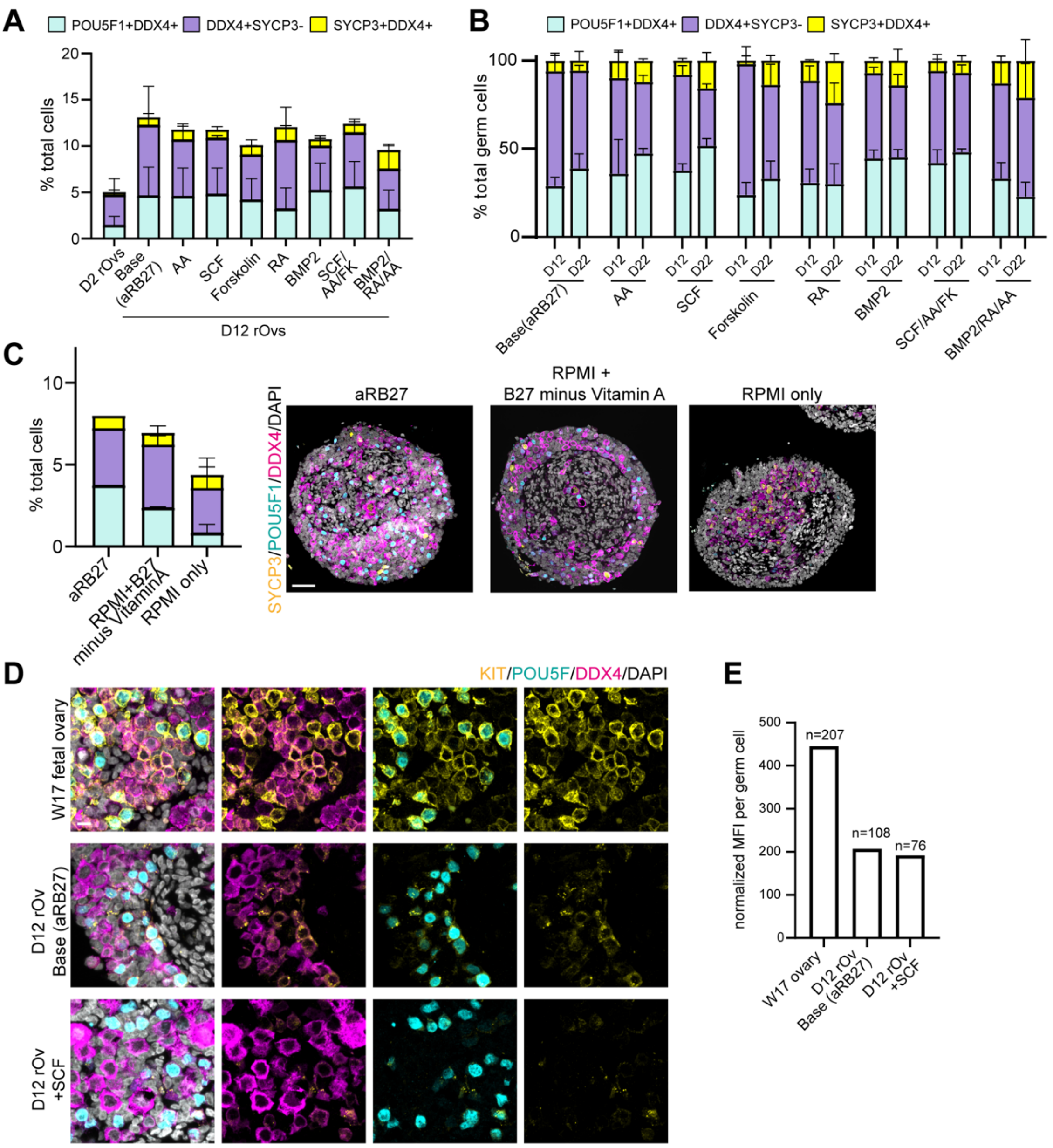


**Supplementary Figure S2. Culturing rOv for up to 22 days.**

(A) Stacked column representation of average percentages of POU5F1+ cells, DDX4+SYCP3- cells and SYCP3+ cells from Figure 2A normalized to total number of germ cells in indicated conditions, pooled from 4 samples (W16, W17, W18, W19), error bar represents SD.

(B) Normalizing the number of POU5F1+ cells, DDX4+SYCP3- cells and SYCP3+ cells from Figure 2D against total number of germ cells, error bar represents SD.

(C) Left: Average percentages of POU5F1+ cells, DDX4+SYCP3- cells and SYCP3+ cells normalized to total number of cells in indicated conditions. 1-4 rOvs per conditions were quantified, error bars represent SD. Right: IF images comparing number of germ cells in indicated conditions. Scale bar: 50 μm.

(D) IF images showing expression of KIT, POU5F1 and DDX4 in both W17 fetal ovary and reconstituted fetal rOvs. Scale bar: 10μm.

(E) Quantification of fluorescence intensity expressed normalized to number of germ cell. (W17 ovary n=207, aRB27 n=108, SCF n=76) N represents number of cells counted.

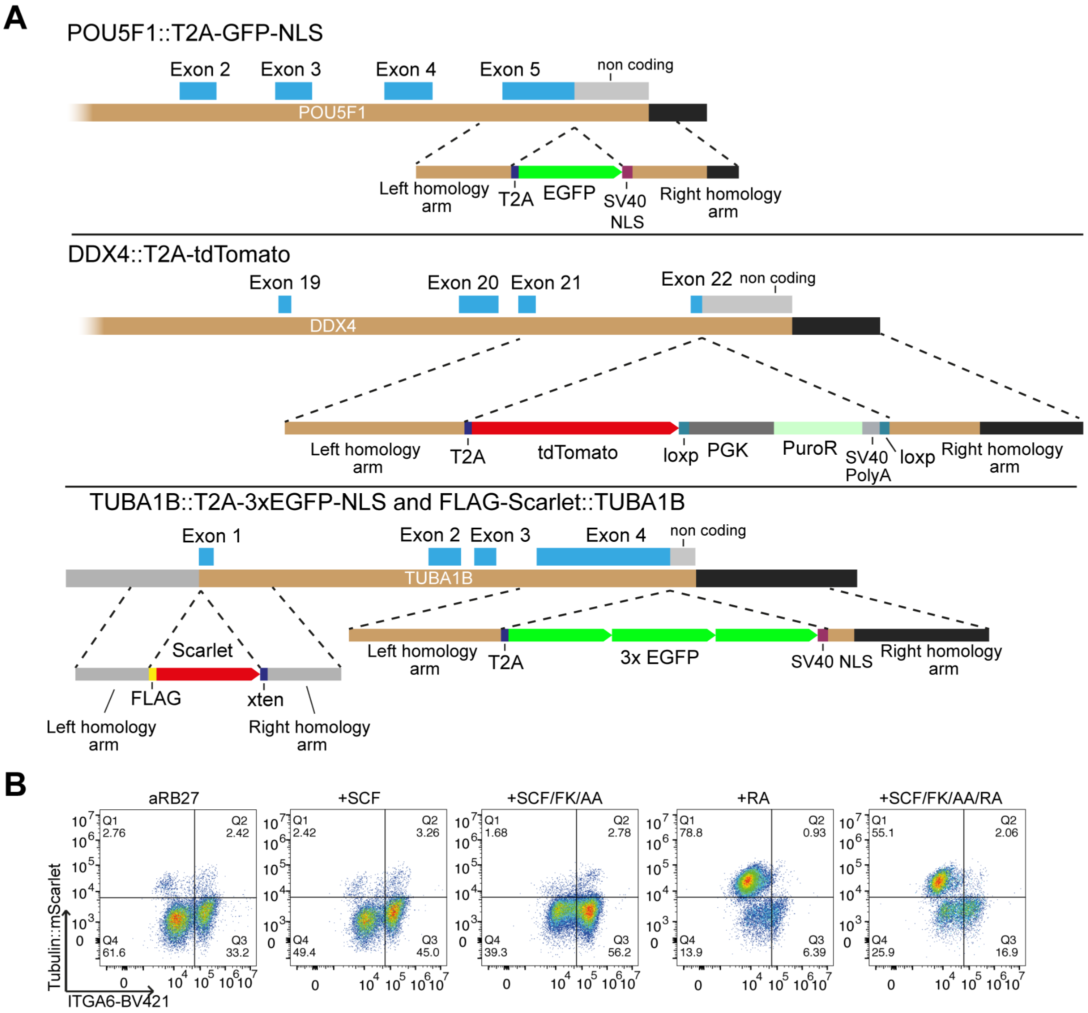


**Supplemental Figure 3. Generation of endogenously tagged hiPSC lines for hPGCLC fetal ovary reconstitution.**

(A) Schematic overviews of endogenous gene tagging knock-ins.

(B) FACS plot showing the percentage of TUBA1B::mSCARLET+ ITGA6+ hPGCLCs in reconstituted rOv.

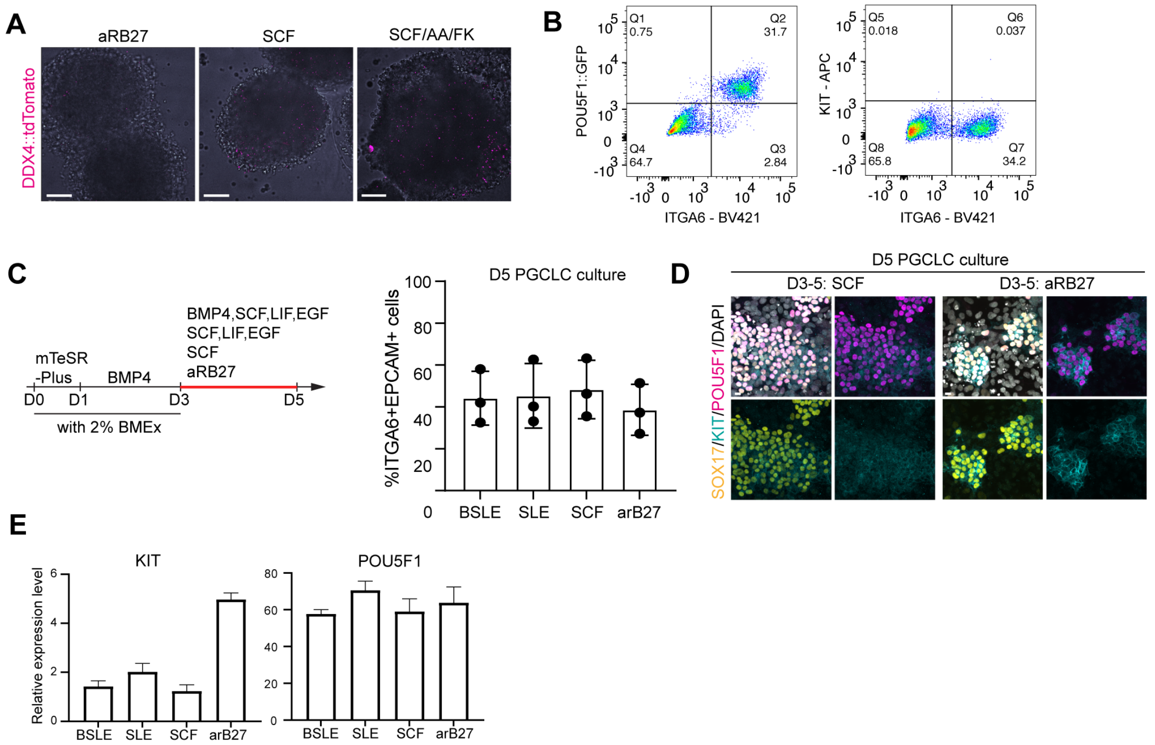


**Supplemental Figure 4. PGCLCs express KIT when SCF is omitted.**

(A) 3D live-cell imaging showing the absence of DDX4::tdTomato signal in the PGCLC/AMLC aggregates in Figure 4E, Scale bar: 100µm

(B) FACS plots showing the expression of POU5F1::GFP, ITGA6, and KIT on D5C17 PGCLCs.

(C) Left: schematics of the PGCLC differentiation protocol with varying D3-5 treatment. Right: D5 PGCLC differentiation efficiency of the from different D3-5 treatment. Dots represent experimental replicate, error bars depict SD. BSLE: BMP4, SCF, LIF, EGF; SLE: SCF, LIF, EGF

(D) Immunofluorescence showing the upregulation of KIT in aRB27 condition compared to SCF, Scale bar: 10µm

(E) qPCR analysis of relative expression levels of *KIT* and *POU5F1*, normalized to *HPRT1*, in D5 differentiation culture for different D3-5 treatment. n=3 technical replicates, error bars represent SD

**Appendix Table S1: Antibodies used in this study**

| **Primary antibodies** | **Host** | **Source** | **Cat. Number** | **Dilution** |
|---|---|---|---|---|
| Oct-3/4(C-10) (POU5F1) | Mouse | Santa Cruz | sc-5279 | 1:200 |
| VASA (DDX4) | Goat | R&D Systems | AF2030 | 1:1000 |
| VASA (DDX4) | Rabbit | Abcam | ab13840 | 1:500 |
| SYCP3 (SCP3) | Goat | R&D Systems | AF3750 | 1:500 |
| HORMAD1 | Rabbit | Abcam | ab220021 | 1:200 |
| ZP3 | Rabbit | Sigma-Aldrich | HPA054061 | 1:200 |
| p63 (TP63) | Mouse | Abcam | ab735 | 1:200 |
| Ki67 | Rabbit | Abcam | ab15580 | 1:500 |
| cKIT (CD117) | Rabbit | DAKO | A4502 | 1:200 |
| FOXL2 | Goat | Novus Biologicals | NB100-1277SS | 1:200 |
| COUP-TFII (NR2F2) | Rabbit | Cell Signalling | ab211776 | 1:200 |
| COL4 | Mouse | Abcam | ab6311 | 1:200 |
| GFP | Chicken | Abcam | ab13970 | 1:500 |
| SOX17 | Goat | R&D Systems | AF1924 | 1:500 |
| DAZL | Rabbit | Abcam | ab215718 | 1:500 |
| BV421 anti-CD49f (ITGA6) | Rat | Biolegend | 313604 | 1:100 |
| BV786 anti-Alkaline Phosphatase (TNAP) | Mouse | Biolegend | 742714 | 1:100 |
| PE/Cyanine7 anti-CD326 (EpCAM) | Mouse | Biolegend | 324222 | 1:100 |
| BV785 anti-CD38 | Mouse | Biolegend | 303529 | 1:100 |
| VioBright FITC anti-SUSD2 | Mouse | Miltenyi | 130-127-902 | 1:100 |
| APC anti-CD117 (KIT) | Mouse | BD Biosciences | 550412 | 1:100 |
| **Secondary antibodies** | | | | |
| Alexa Flour 488 donkey anti-goat IgG | Donkey | Thermo Fisher Scientific | A-11055 | 1:500 |
| Alexa Flour 647 donkey anti-rabbit IgG | Donkey | Thermo Fisher Scientific | A-31573 | 1:500 |
| Alexa Flour 647 donkey anti-mouse IgG | Donkey | Thermo Fisher Scientific | A-31571 | 1:500 |
| Alexa Flour 488 donkey anti-mouse IgG | Donkey | Thermo Fisher Scientific | A-21202 | 1:500 |
| Alexa Flour 555 donkey anti-rabbit IgG | Donkey | Thermo Fisher Scientific | A-31572 | 1:500 |
| Alexa Flour 647 donkey anti-goat IgG | Donkey | Thermo Fisher Scientific | A-21447 | 1:500 |
| Alexa Fluor 555 donkey anti-chicken IgY | Donkey | Thermo Fisher Scientific | A78949 | 1:500 |

**Appendix Table S2: List of primers**

| Homology arm cloning primers | | |
|---|---|---|
| Target | Forward (5' - 3') | Reverse (5' - 3') |
| DDX4-right-arm | GTTTTGATGCAGAGAAGAAAATAGTTTTG | GGAAACAGCTATGACCgTGTCTCCCCAGTATTTACACCTCAC |
| DDX4-left-arm | TGTAAAACGACGGCCAGTGGGCTCAACAGGATGTTCCTGCAT | AGAAGACTTCCCCTGCCCTCATCCCATGACTCATCATCTACTGGATTG |
| TUBA1B-right-arm | GGCACTGCAGCATGTCATGCTCCCAG | GCAGCCATAATTTTCCTGTGCTTTCCG |
| TUBA1B-left-arm | TCTGTTTTGCCAATTCCTTGTGC | TTAGTATTCCTCTCCTTCTTCCTCACCCTC |
| gRNAs | | |
| Name | Sequence (5’-3’) | |
| TUBA1B N-terminus | GATGCACTCACGCTGCGGGA | |
| TUBA1B C-terminus gRNA1 | TGACATGCTGCAGGGCCAAA | |
| TUBA1B C-terminus gRNA2 | TTTGGCCCTGCAGCATGTCA | |
| DDX4 gRNA1 | CATCAAAACCACAGACTTGA | |
| DDX4 gRNA2 | CAAAACATCCTTCAAGTCTG | |
| AX33_pgRNAOCT4.1 | GCACCTCAGTTTGAATGCAT | |
| real-time PCR primers | | |
| Target | Forward (5' - 3') | Reverse (5' - 3') |
| *POU5F1* | CATCAAAGCTCTGCAGAAAGAACT | CTGAATACCTTCCCAAATAGAACCC |
| *NANOS3* | CAAGGCGAAGACACAGGACA | TCCTAGGTGGACATGGAGGG |
| *KIT* | GATGGATGGATGGTGGACAC | GGGATTTTCTCTGCGTTCTG |
| *DPPA3* | TAGCGAATCTGTTTCCCCTCT | CTGCTGTAAAGCCACTCATCTT |
| *DPPA5* | CACCGAGGTCGTGGTTTACG | GGCCTAGTTCGAGGGCATTC |
| *CDH5* | CGGCGCCAAAAGAGAGATTG | CGGAAGACCTTGCCCACATA |
| *SMARCA2* | TCCTCGCGAGCAAGCATTA | ATCATGCTGTGGACGGAACC |
| *ESAM* | ATGTGACGCTGGAAGTGAGC | AGCAATGGCATCCTCCTTGATA |
| *GAPDH* | AAGGTGAGGGTCGGAGTCAAC | GGGGTCATTGATGCCAACAATA |
| *HPRT1* | TGACACTGGCAAAACAATGCA | GGTCCTTTTCACCAGCAAGCT |
